# Clumpy coexistence in phytoplankton: The role of functional similarity in community assembly

**DOI:** 10.1101/869966

**Authors:** Caio Graco-Roza, Angel M Segura, Carla Kruk, Patrícia Domingos, Janne Soininen, Marcelo Manzi Marinho

## Abstract

Emergent neutrality (EN) suggests that species must be sufficiently similar or sufficiently different in their niches to avoid interspecific competition. Such a scenario results in a transient pattern with clumps and gaps of species abundance along the niche axis (e.g., represented by body size). From this perspective, clumps are groups of coexisting species with negligible fitness differences and stochastic abundance fluctuations. Plankton is an excellent model system for developing and testing ecological theories, especially those related to size structure and species coexistence. We tested EN predictions using the phytoplankton community along the course of a tropical river considering (i) body size structure, (ii) functional clustering of species in terms of morphology-based functional groups (MBFG), and (iii) the functional similarity among species concerning their functional traits. Two main clumps in the body size axis (clump I and II) were conspicuous through time and were detected in different stretches of the river. Clump I comprised medium-sized species from the MBFGs IV, V, and VI while clump II included large-bodied species from the MBFGs V and VI. Pairwise differences in species biovolume correlated with species functional similarity when the whole species pool was considered, but not among species within the same clump. Although clumps comprised multiple MBFGs, the dominant species within the clump belonged always to the same MBFG. Also, within-clump species biovolume increased with functional distinctiveness considering both seasons and stretches, except the lower course. These results suggest that species within clumps behave in a quasi-neutral state, but even minor shifts in trait composition may affect species biovolume. Our findings point that EN belongs to the plausible mechanisms explaining community assembly in river ecosystems.

## Introduction

Understanding the mechanisms promoting species coexistence and shaping community structure has been a long-standing goal in community ecology. The former idea that the number of coexisting species is limited by the number of growth-limiting resources or niche dimensions (Gause 1936, Hardin 1960) and its derivate idea, “*the paradox of the plankton”* (Hutchinson 1957), have been widely explained in terms of endogenous and exogenous spatio-temporal mechanisms (Roy and Chattopadhyay 2007). Trait-based approaches are useful to test this matter due to their potential to generalize patterns beyond species’ identity, especially because traits influence the species’ ability to acquire resources and persist through environmental changes (McGill et al. 2006, Díaz et al. 2013, 2016). Nonetheless, the niche-based theory proposes that the environment filters community composition through species’ ecological requirements, which can be perceived through species’ traits. Also, intra- and inter-specific interactions potentially drive community assembly, in local communities (Götzenberger et al. 2012). In contrast, the more recent neutral theory suggests that diversity results from random dispersal, speciation, and extinction rates with no role of niche differences in species coexistence (Hubbell 2001). This type of dynamics should then result in a random distribution of functional traits along environmental gradients (Kraft et al. 2008, Cornwell and Ackerly 2009).

More recently, it was shown that community organization is driven by eco-evolutionary processes such as speciation and nutrient uptake kinetics resulting in groups comprising different species with similar ecological requirements (Gravel et al. 2006, Scheffer and van Nes 2006, Hubbell 2006). This finding led to the ‘emergent neutrality hypothesis’ (EN; Holt, 2006) that has been supported by observational studies, e.g., for phytoplankton from brackish waters (Segura et al. 2011), birds from the North of Mexico (Thibault et al. 2011) and beetles at the global scale (Scheffer et al. 2015). EN suggests that species must be sufficiently similar, and thus, behave neutrally, or different enough in their niches to avoid competition. Such a scenario would result in species-rich aggregations or clumps along the niche axis (Scheffer and van Nes 2006, Vergnon et al. 2009, Fort et al. 2010). Modelling studies have shown that such predictions apply for both steady environmental conditions (Fort et al. 2010), and also fluctuating resource conditions (Sakavara et al. 2018). Empirical evidence about EN is still scarce, however (Scheffer et al. 2018).

The clumpy pattern arises from the exceedingly slow displacement rate of species under intense competition, that is, species within the same clump overlap in their niche such that the displacement rate of competing species is similar to the competition at the intraspecific level, leading to stochastic fluctuations in species abundances through time (Scheffer et al. 2018). Thus, the number of clumps corresponds to the number of species to be expected to stably coexist at equilibrium, but the identity of the dominant species is expected to be random among the clump residents. However, the assignment of species to clumps is challenged by the fact that trait differences among species are continuous (Villéger et al. 2008) and the threshold to include a species within a clump varies with the statistical approach that is applied (Segura et al. 2011, D’Andrea et al. 2019). Therefore, given this methodological limitation, it is difficult to state empirically whether species behave neutrally within clumps (i.e., when the strength of interspecific interactions equals the intraspecific interactions) or if results are an artefact of clump construction.

Zooming in on the uniqueness of trait combinations of species, i.e., functional distinctiveness, within clumps may advance our comprehension of biotic interactions and move towards a measurable value of similarity at which species coexistence is driven stochastically. Functional distinctiveness reflects the non-shared functions among species within a given species pool (Violle et al. 2017), mirroring the concept of functional similarity (Pavoine et al. 2017). However, functional distinctiveness is not directly linked to functional similarity at the pairwise level (Coux et al. 2016, Ricotta et al. 2016, Violle et al. 2017). For example, two species may be equally distinct, i.e. the degree to which a species differs from all the others within the species pool concerning their functional traits, and still not be similar in their trait composition at a pairwise level (Coux et al. 2016). This suggests that both pairwise functional similarity and group-based functional distinctiveness are complementary metrics to assess the role of trait combination in community assembly. To this end, phytoplankton communities are useful for biodiversity theory testing due to their species-rich communities, rapid responses (in human time-scales) and well-characterized relationships between morphology and physiological and ecological responses (Litchman and Klausmeier 2008, Kruk and Segura 2012, Litchman et al. 2012).

Body size is considered a master ecological trait and it is often used to characterize species niche differences (Downing et al. 2014). In phytoplankton, the body size is related to physiology and life-history (Litchman and Klausmeier 2008), photosynthetic processes (Marañón 2008), nutrient uptake kinetics (Litchman et al. 2010) and other eco-evolutionary processes, e.g. the relationship among predation rates, nutrient uptake and organisms body size (Sauterey et al. 2017). Although body size may relate to different processes, using a single trait as a proxy for niche differences may not evidence species differences generated by hidden/unknown niche axes (i.e. ecological dimensions of the niche) and impair the understanding of clumpy patterns (Barabás et al. 2013, D’Andrea et al. 2018). The use of multiple traits emerges as a powerful tool to disentangle phytoplankton functional structure and evaluate competing hypotheses (Reynolds et al. 2014, Chen et al. 2015, Bortolini and Bueno 2017, Aquino et al. 2018). Morphology-based functional groups (MBFG) classification of phytoplankton species (Kruk et al. 2010) is a multidimensional combination of morphological traits that cluster organisms into seven groups with similar physiology and ecological responses, potentially overcoming the limitations of using a single trait dimension only. Assessing the functional distinctiveness of species within the same functional cluster (e.g., clumps, MBFGs) could help to study the existence of functional equivalence (i.e., neutrality) among species. Overall, the functional similarity among species is a useful tool to compare species in a multidimensional space, particularly because the environment may filter different functional traits across space and time (D’Andrea et al. 2020).

Rivers are highly heterogeneous systems characterized by a continuous water flow that affects the ecosystem’s morphology (e.g., meandering), sedimentation patterns, organisms’ dispersal, and more specifically the phytoplankton abundance and distribution (Reynolds and Descy 1996, Wetzel 2001). Several theories, e.g., the River Continuum (Vannote et al. 1980) and Flood Pulse (Junk et al. 1989) concepts explain the longitudinal distribution and abundance of riverine phytoplankton communities. However, an explicit study of communities’ body size structure and species coexistence under EN in riverine ecosystems is lacking. For example, phytoplankton species should attain higher biomass at the middle reaches or in the upper reaches of low-gradient stretches (Descy et al. 2017). Also, competition rates vary along the river course because water turbulence reduces the likelihood of biotic interactions (Reynolds et al. 1994), meaning that clumpy coexistence may not be observed in riverine phytoplankton. Alternatively, if functional trait combinations of species within the local species pool result from eco-evolutionary processes (Scheffer et al. 2015), the clumpy pattern should also be apparent in riverine phytoplankton communities. Here, we push forward three hypotheses to be tested in a tropical river by investigating phytoplankton community size structure both seasonally and spatially. We expect that:

**H**_**1**_ – There are peak aggregations of species abundance (i.e., clumps) along the body size axis of phytoplankton in the river that remain constant across space and time as a result of eco-evolutionary processes.

**H**_**2**_ – Pairwise-differences in species abundances increase with functional dissimilarity at the community-level but not at the clump level because species within the same clump behave in a quasineutral state. Thus, the dominance within clump varies stochastically between species as fitness differences are negligible.

**H**_**3**_ – Species abundance increases with functional distinctiveness with respect to other species within the clumps. Although abundance fluctuates stochastically at the pairwise level, the number of species bearing similar trait combinations may affect the likelihood of the interactions within clumps. Therefore, species with the most distinct trait combinations concerning their clump peers are less likely to share the same ecological requirements, and by consequence, attain higher abundance.

## Methods

### Study area

Samples were taken monthly at nine stations along the Piabanha river between May 2012 and April 2013. Piabanha river is in the tropical region of Brazil and has a drainage basin of approximately 4500 km^2^ (Figure 1). The headwater is on Petrópolis at 1546m altitude and drains to the medium valley of Paraíba do Sul River crossing three cities and with agricultural activities in their watershed. We set three river stretches (lower, medium, and upper courses) based on the location of steep slopes on the river elevation profile (Figure 1). Data from two meteorological stations (Bingen and Posse; Figure 1), located in the upper and lower courses of the river, were used to measure rainfall. We analysed meteorological data up to three days before each sampling campaign. We then classified seasons as a dry season (May - October) and a wet season (November – April) based on the rainfall data.

**Figure 1.**
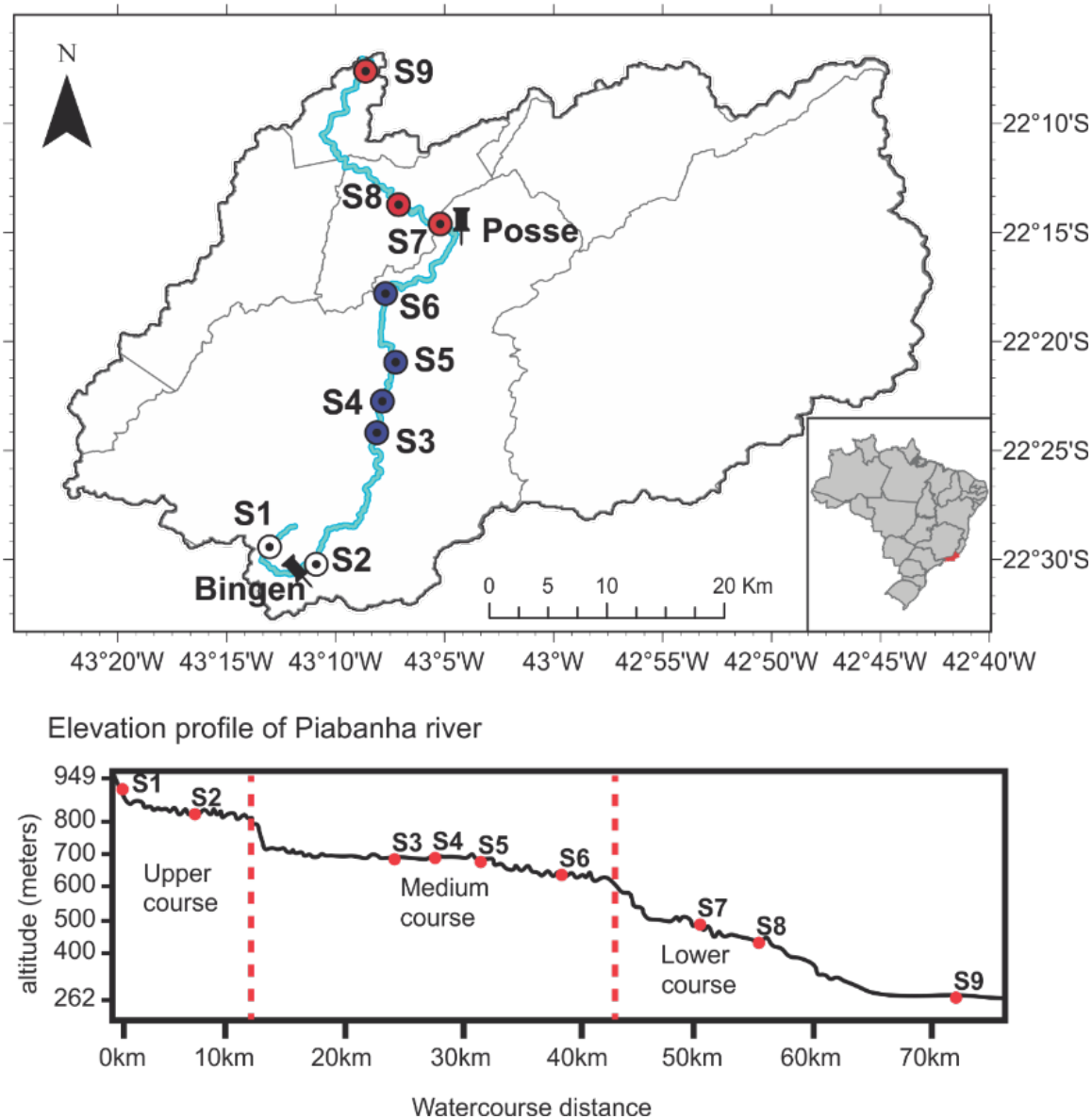
Map of the study area. The watershed area of the Piabanha river showing the river course (blue line), the meteorological stations Bingen and Posse, and the sampling sites are coloured according to river stretches (white circles = upper course, blue circles = medium course, red circles = lower course). The vertical dotted red line in the elevational profile figure indicates the locations of steep slopes used to define the boundaries of the river stretches.

### Sampling and sample analysis

In the field, we measured water temperature (°C), dissolved oxygen (DO, mg L^− 1^), and turbidity by a multiparameter probe sonde (YSI model 600 QS). Water discharge (WD, m^3^s^− 1^) was measured with the SonTek RiverSurveyor – M9. Furthermore, water samples were taken and kept frozen (one or 2 weeks) until the laboratory analysis for ammonium (N·NH4+, mg L^− 1^), nitrate (N·NO_3_ ^−^, mg L^− 1^), nitrite (N·NO_2_ ^−^, mg L^− 1^), total phosphorus (TP, mg L^− 1^) and soluble reactive phosphorus (SRP, mg L^− 1^) (Figure 2). Ammonium, nitrite, and nitrate were summed up and are expressed as dissolved inorganic nitrogen (DIN, mg L^− 1^). The water samples were filtered (except for total phosphorus analysis) using borosilicate filters (Whatman GF/C), and nutrient concentrations were measured following APHA (2005). A complete description of the spatial and seasonal patterns of the environmental variables measured in the Piabanha river can be found in Graco-Roza et al. (2020).

**Figure 2.**
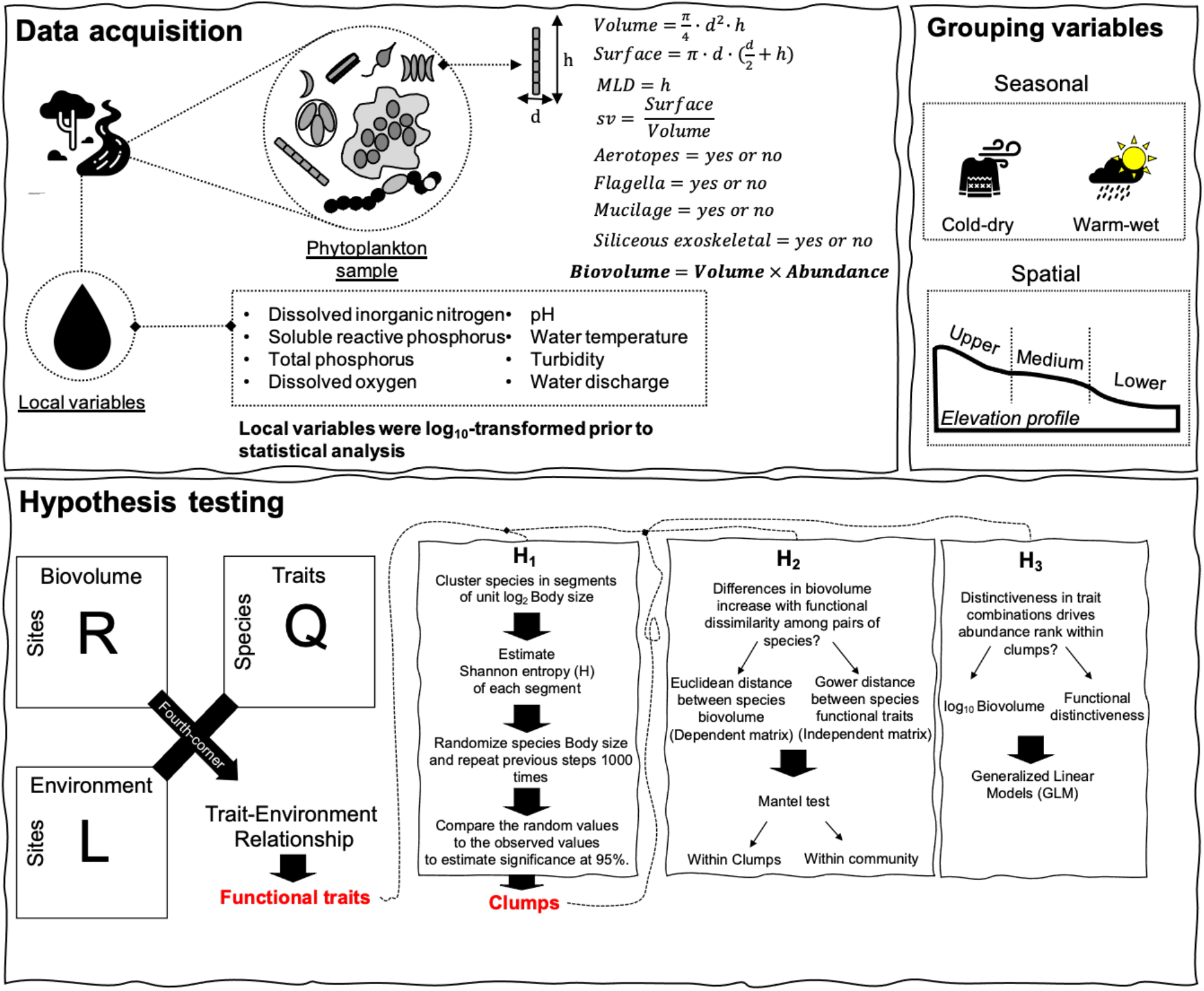
Study design. Sketch diagram showing the steps from data acquisition to hypothesis testing, and the grouping variables used to analyse the data. Water samples were taken from the river for the estimation of local environmental variables, and phytoplankton qualitative and quantitative analysis. Phytoplankton species were measured based on approximate geometrical forms for the estimation of individual volume and surface. Other traits were also measured during the quantitative and qualitative analysis. Samples were divided into seasonal and spatial groups prior to the analysis. Species biovolume were used along with its traits to test the relationship between trait and environment, and to highlight the functional traits. Only body size was used to test the presence of significant clumps (H_1_), while the other functional traits were used to test the effects of functional similarity within significant clumps (H_2_-H_3_).

### Phytoplankton samples

Subsurface samples of phytoplankton were collected with a bottle of 200 mL and fixed with Lugol. In the laboratory, phytoplankton species were identified, and population densities were estimated under an inverted microscope (Olympus CKX41) (Utermöhl 1958). At least 100 individuals of the most abundant species were counted in each sample (Lund et al. 1958, Uhelingher 1964). Biovolume (mm^3^L^− 1^) of phytoplankton species was estimated by multiplying the density of each population (ind. L^-1^) by the average individual volume of the species (V, µm^3^org^-1^). The volume of each species was estimated by measuring geometrical dimensions and approximating to defined geometrical forms following Hillebrand et al. (1999). Geometrical dimensions were measured in 20 organisms from each species (when possible) and the average was used to characterize individual bod size (volume). We recall that biovolume represents the biomass density and volume is an organism’s trait. Species’ surface area (S, µm^2^) was estimated, the maximum linear dimension (MLD, µm) was measured, and the presence or absence of aerotopes, mucilage, flagella, and siliceous exoskeletal structures was noted in each species (Figure 2). We then used the volume and surface area of the species to estimate the individual surface volume ratio (SV). Species were then classified into MBFG according to Kruk et al. (2010), based on the above mentioned morphological traits. This classification included the following seven groups: (I) small organisms with high SV, (II) small, flagellated organisms with siliceous exoskeletal structures, (III) large filaments with aerotopes, (IV) organisms of medium size lacking specialised traits, (V) flagellates unicells with medium to large size, (VI) non-flagellated organisms with siliceous exoskeletons and (VII) large mucilaginous colonies. For further details on MBFG classification, we refer to Kruk et al. (2010) or Segura et al. (2013a). The information on the traits measured for each species, and the classification into MBFGs can be found in the Table S1.

### Traits-environment relationship

We tested the relationship between morphological traits and the environmental variables using a three-table ordination (RLQ) combined with a fourth-corner analysis (Dray et al. 2014) (Figure 2). Both RLQ and fourth-corner methods require the information from three tables: (i) a data frame including the measurements of environmental variables across the sampling sites (R table), (ii) a matrix containing species abundances or occurrences across the sampling sites (L table), and (iii) a data frame comprising the trait values for each species (Q table). Also, both methods rely on the analysis of the fourth-corner matrix, crossing the information between tables R and Q, weighted by table L. The RLQ analysis (Legendre et al. 1997) provides ordination scores to summarize the joint structure among the three tables, but it does not allow the identification of traits or environmental variables contributing significantly to the structure. The fourth corner method (Dolédec et al. 1996) tests the significance of bivariate associations between each trait and environmental variables but disregards the covariance among traits or environmental variables. Here, we combined the RLQ analysis with the fourth corner method by applying the fourth corner method to the output of the RLQ analysis instead of the original raw values (Dray et al. 2014). By doing this, we summarized the main patterns in the multivariate space and tested the global significance of the trait–environment relationships using the *S*_RLQ_ multivariate statistic and the fourth corner sequential testing procedure (Dray and Legendre 2008). Applying the fourth corner method in the output of the RLQ determines (i) the relationship between individual traits and RLQ environmental scores (a.k.a. environmental gradients, and (ii) the relationship between environmental variables and RLQ traits scores (a.k.a. trait syndromes)(Dray et al. 2014).

Before applying the RLQ method, we first log-transformed (log_10_ x+1) species biovolume, species traits (SV, MLD, and V), and environmental variables (except pH and temperature). A correspondence analysis (Benzécri 1973) was performed on the L table using the function dudi.ca from ‘ade4’, and a Hill-Smith analysis (Hill and Smith 1976) on the R and Q tables separately using the function dudi.hill from ‘ade4’. We used Hill-smith analysis because both R and Q table included categorical or binary variables. The RLQ analysis was conducted in the output of the ordinations using the function rlq from ‘ade4’. We tested the significance of the joint structure among the RLQ tables using the fourth-corner method with a stepwise permutation procedure of 999 permutations using the function rlq.fourthcorner from ‘ade4’. The null hypothesis that given fixed traits, species abundances are independent of environmental conditions was evaluated by permuting sites (rows of tables L or R) while keeping the species traits (table Q) fixed. The null hypothesis that given fixed environmental conditions, species abundances are independent of functional traits was evaluated by permuting species (columns of table L or rows of table Q) while the environmental conditions (R table) were kept fixed (Dray et al. 2014). Rejecting both null hypotheses imply that tables R, L, and Q are significantly linked. Because the fourth-corner analysis explores one trait and one environmental variable at a time, multiple statistical tests are performed simultaneously increasing the probability of type I error (i.e. false significant associations), thus we adjusted p-values for multiple testing using the false discovery rate method (Benjamini and Hochberg 1995). We divided the value of the fourth-corner correlation by the square-root of the first eigenvalue of the correspondence analysis of the L matrix, which is the maximum possible value (Peres-Neto et al. 2017).

### Clumpy patterns

To test for the existence of peak aggregations of species biovolume along the body size axis of phytoplankton - **H**_**1**,_we analysed the community structure in each season (dry and wet) and river stretches (upper, medium, and lower course) (Figure 2). First, the individual volume of species was log-transformed (log_2_) and used as the main niche axis (X= log_2_ volume) following Segura et al. 2011. Hence, we divided the niche axis into equally spaced segments (one segment per unit log_2_ volume) and for each segment (j), we estimated the Shannon entropy (H) using the biovolume of the observed species (Fort et al. 2010, Segura et al. 2011). The entropy index was defined as:

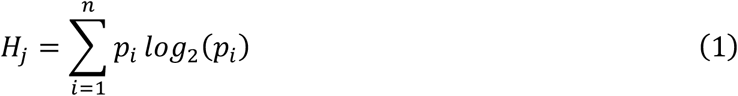

here *p*_*i*_ is the fraction of biovolume of species *i* in the community of *n* species. Finally, we tested the significance of the entropy (*H*) by comparing the observed *H* against an expected uniform distribution under the null hypothesis of homogeneous *H*. For this, we created 1000 communities by sampling the volume of species from a random uniform distribution bounded by observed individual volumes. Then, each species had a biovolume assigned to it, which was taken from randomization of the observed biovolume matrix, keeping both the empirical species rank-biovolume pattern and total biovolume in the sample. For each segment, the observed *H* was compared with the distributions of *H* generated under the null hypothesis, with significance defined according to standard 5% criterion (Fort et al. 2010, Segura et al. 2011). Finally, we considered a significant segment or two consecutive significant segments as a clump.

### Functional dissimilarity

To test whether differences in species biovolume increases with functional dissimilarity – **H**_**2**_, we first calculated the functional dissimilarity and the differences in biovolume among pairs of species using the whole community and using only the species from the significant clumps separately (Figure 2). The functional dissimilarity was obtained by calculating Gower’s general dissimilarity coefficient on all the species functional traits, that is, all the traits that showed a significant (p < 0.05) relationship with the environmental gradients in the fourth corner method. The dissimilarity coefficient was estimated using the function gowdis from ‘FD’. We used Gower’s dissimilarity (*Gd*) because it can handle mixed variable types (continuous, binary, ordinal, and categorical traits). *Gd* defines a distance value *d*_*jk*_ between two species as:

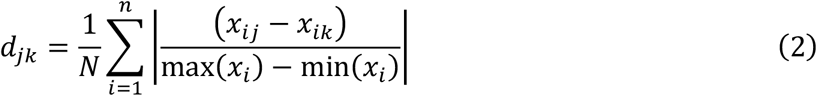

where, *N* is the number of functional traits considered, *x*_*ij*_ the value of trait *i* for species *j*, and *x*_*ik*_ the value of the trait *i* for species *k*. Therefore, *Gd* = d*jk* = functional dissimilarity. We thus tested **H**_**2**_ by conducting Mantel tests with 999 randomizations on the matrices of functional dissimilarity and differences in biovolume using the function mantel from ‘vegan’. We performed the Mantel test considering: (i) all species present in a given season or river stretch, and (ii) separately for the species of each significant clump that were present in a given season or river stretch.

### Functional distinctiveness (F_Dist_)

To test whether species biovolume increases with functional distinctiveness at the clump-level – **H**_**3**_, we estimated the functional distinctiveness (F_Dist_) as the Euclidean distance of a species to the average trait position (centroid) in the multidimensional functional space for the set of species of each of the significant clumps using the equations proposed by Anderson (2006) (Figure 2). First, we applied a Principal Coordinates Analysis (PCoA) in the species-by-traits data table using Gower’s dissimilarity (*Gd*) and obtained species coordinates in the functional space using all the axes from the PCoA. Hence, F_Dist_ was calculated as:

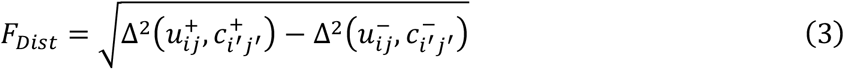

where Δ^2^ is the squared Euclidean distance between u_ij_, the principal coordinate for the *j*th species in the *i*th clump, and c_i_, the coordinate of the centroid for the *i*th clump. The super-scripted ‘+’ and ‘-’ indicate the real and imaginary parts respectively (see Anderson 2006, for details). We did not weight the clump-centroid by species biovolume because it would artificially give higher distinctiveness for less abundant species and bias our analysis. Besides, we calculated F_Dist_ using only species from the significant clumps and normalized the F_Dist_ to range between zero and one by dividing the actual F_Dist_ values by the F_Dist_ of the most distinct species of the clump. We tested the **H**_**3**_, by modelling the relationship between species biovolume and F_Dist_ using linear models. We used log_10_ biovolume as the dependent variable, with F_Dist_ and Clump (i.e., the clump to which a species belong) as the independent variables for each season and river stretch separately.

### Statistical analyses

Statistical analyses were performed on R v.4.0.4 (R Core Team 2020) using the packages ‘ade4’ v.1.7.16 (Chessel et al. 2004, Dray and Dufour 2007, Dray et al. 2007, Bougeard and Dray 2018, Thioulouse et al. 2018), ‘FD’ v.1.0.12 (Laliberte et al. 2010, Laliberté et al. 2014), the suite of packages ‘tidyverse’ v.1.3.0 (Wickham et al. 2019), and the package ‘vegan’ v.2.5.7 (Oksanen et al. 2020). The code used to generate results can be found at https://github.com/graco-roza/clumpy-coexistence-phytoplankton.

## Results

Our samples included 150 species that were classified in six (MBFG I, III, IV, V, VI, and VII) from the seven MBFGs based on their functional traits (Table 1). MBFGs IV, V, and VI included 87% of the total number of species. Species from MBFG IV included filamentous, colonial, and unicellular species ranging from 21 µm^3^ to 8181 µm^3^ lacking specialized morphological traits (e.g., flagella, siliceous exoskeletal structures). MBFG V comprised unicellular flagellated species ranging in volume from 31µm^3^ to 31864 µm^3^, and MBFG VI included unicellular and chain-forming species with a siliceous exoskeletal body that ranged in volume from 48µm^3^ to 19045µm^3^.

**Table 1.** Distribution of species among the morphological-based functional groups.

| MBFG | Number of species | Representative taxa |
| --- | --- | --- |
| I | 9 | *Chroococcales* sp., *Chroococcus* sp. |
| III | 3 | *Limnothrix* sp. |
| IV | 60 | *Pseudanabaena limnetica*, *Pseudanabaena catenata* |
| V | 13 | *Euglena* sp., *Cryptomonas* sp. |
| VI | 57 | *Cymbella* sp., *Synedra* sp. |
| VII | 8 | *Dictyosphaerium* sp. |
| Total | 150 |  |

Regarding the trait-environment relationship, the first two RLQ axes preserved well the variance of the ordinations and explained altogether 84.37 % of the variation, with 66.71% corresponding to the first axis alone. The *S*_RLQ_ statistic indicated a significant global relationship between trait syndromes and environmental gradients (*r* = 0.18, p-value < 0.01). Mainly, the first trait syndrome (RLQ axis 1_Trait_) correlated significantly with the first environmental gradient (RLQ axis 1_Environment_; *r* = 0.17, p < 0.01) while the second trait syndrome (RLQ axis 2_Trait_) correlated significantly with the second environmental gradient (RLQ axis 2_Environment_; *r* = 0.12, p < 0.01). Yet, there was no significant relationship between the first trait syndrome and the second environmental gradient, nor between the second trait syndrome and the first environmental gradient (Table 2).

**Table 2.** Combined fourth-corner-RLQ analysis to test the relationship between functional syndromes (RLQ axis_Trait_) and environmental gradients (RLQ axis_Environment_).

|  | RLQ axis 1<sub>Trait</sub> | RLQ axis 2<sub>Trait</sub> |
| --- | --- | --- |
| | Pearson's $r$ (adjusted $p$-value) | |
| RLQ axis 1<sub>Environment</sub> | **0.17 (< 0.01)** | 0.00 (1.00) |
| RLQ axis 2<sub>Environment</sub> | 0.00 (1.00) | **0.12 (< 0.01)** |

Results of the RLQ showed that most of the biovolume-based variation in trait syndromes were related to the flow regime of the river Piabanha. The first RLQ axis summarised the spatial gradient in the environmental conditions, specifically the increase in turbidity, temperature, pH, and water discharge from the upper to the lower courses while the second RLQ axis summarised the seasonal gradient with the smaller nutrient concentrations and higher water turbidity in the wet season (Figure 3A). Noteworthy, the spatial and seasonal gradients in abiotic conditions coupled with the distribution of species from different MBFGs. The MBFGs I, III, and IV attained higher abundances in the upper course, contrasting with MBFG VI that showed the highest abundances at the lower course (Figure 3B). Regarding the seasonal gradient, MBFG V had higher abundances during the dry season (Figure 3B). Indeed, the fourth-corner method showed that the first trait syndrome correlated positively with turbidity (*r* = 0.16, p = 0.01), temperature (*r* = 0.14, p = 0.02), pH (*r* = 0.13, p = 0.03), water discharge (*r* = 0.13, p = 0.03) and total phosphorus (*r =* 0.12, p = 0.04), and negatively with the upper course (*r* = -0.21, p < 0.01; Figure 3C). Besides, the second trait syndrome correlated negatively with dissolved inorganic nitrogen (*r* = -0.11, p = 0.04) and positively with turbidity (*r* = 0.13, p = 0.01; Figure 3C). For the spatial environmental gradient, there was a positive correlation with the species volume (*r* = 0.15, p = 0.01) and the presence of siliceous exoskeletal structures (*r* = 0.22, p < 0.01), and a negative correlation with surface volume ratio (*r* = -0.19, p < 0.01; Figure 3D). For the seasonal environmental gradient, there was a positive significant correlation with species maximum linear dimension (*r* = 0.12, p = 0.01), and a negative significant correlation with the presence of flagella (*r* = -0.12, p = 0.01; Figure 3D).

**Figure 3.**
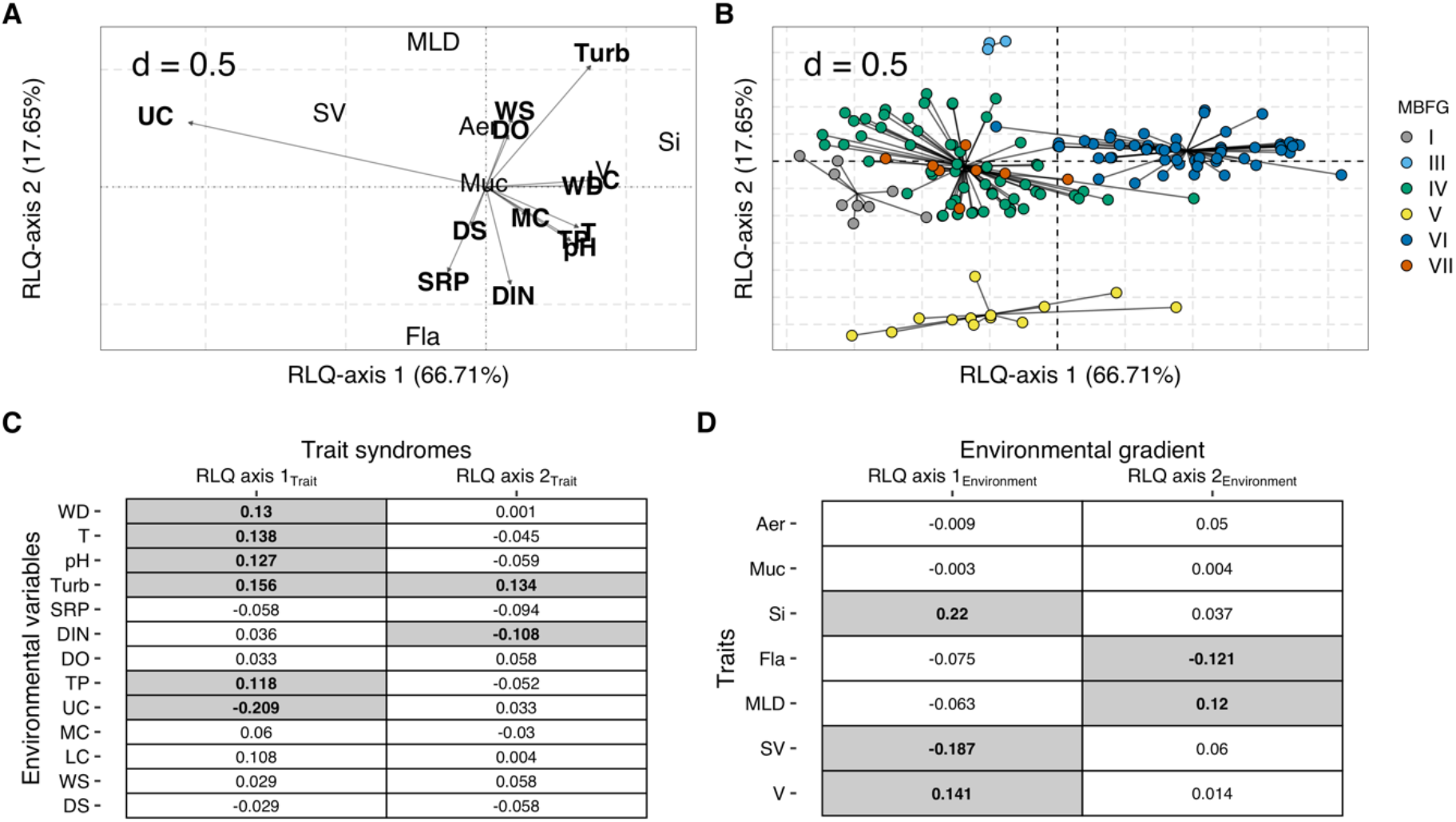
Results of the (A, B) RLQ ordination and (C, D) hypothesis testing through fourth-corner analysis. A) The relationships between species traits and environmental variables. B) The distribution of species in the functional space. Each point in the ordination plot represents the position of a species modelled according to its traits on RLQ axes 1 and 2. The black lines connect the species to the centroid of its morphology-based functional groups - MBFG. Colours represent MBFGs, C) The correlation between species traits and the environmental gradients (RLQ axis _environment_), and D) the relationship between environmental variables and the trait syndromes (RLQ axis _trait_). The grey boxes in C and D indicate significant relationships, and the values within the boxes indicate the Pearson’s *r*. Aer, aerotopes; Muc, mucilage; Si, siliceous exoskeletal structures; Fla, flagella; MLD, maximum linear dimension; SV, surface volume ratio; V, volume; WD, water discharge; T, temperature; Turb, turbidity; SRP, soluble reactive phosphorus; DIN, dissolved inorganic nitrogen; DO, dissolved oxygen; TP, total phosphorus; WS, wet season; DS, dry season; UC, upper course; MC, medium course; LC, lower course.

Overall, species volume ranged from 4.19 µm^3^ to 31864 µm^3^ totalling 14 equally spaced segments (S) of volume along the niche axis. From the 14 segments, three of them showed significant (p < 0.05) entropy values, specifically S9, S13, and S14. This resulted in a biovolume aggregation (i.e., clumps) in two regions of the niche axis considering both seasonal (Figure 4; Column 1) and spatial categories (Figure 5; Column 1). The first clump included 24 species from the MBFGs IV, V, and VI at the range of S9 (512µm^3^ - 1024µm^3^), particularly eight species from MBFG IV (e.g., *Pseudanabaena catenata* and *P. limnetica*), four species from the MBFG V (e.g., *Strombomonas* sp., *cf Cryptomonas* sp.), and 12 species from MBFG VI (e.g., *Fragillaria capuccina* var. gracilis, *Achnantes cf. rupestoides*). The second clump (hereafter Clump II) included six species at the range of S14, being two species from MBFG V (i.e., *Euglena* sp.) and four species from MBFG VI (e.g., *Pinnularia* sp., *Synedra* sp.). However, during the wet season, the S13 also had significant entropy values and six more species from MBFG VI (e.g., *Achnantes inflata, Cymbella* sp.) were included in the clump II.

**Figure 4.**
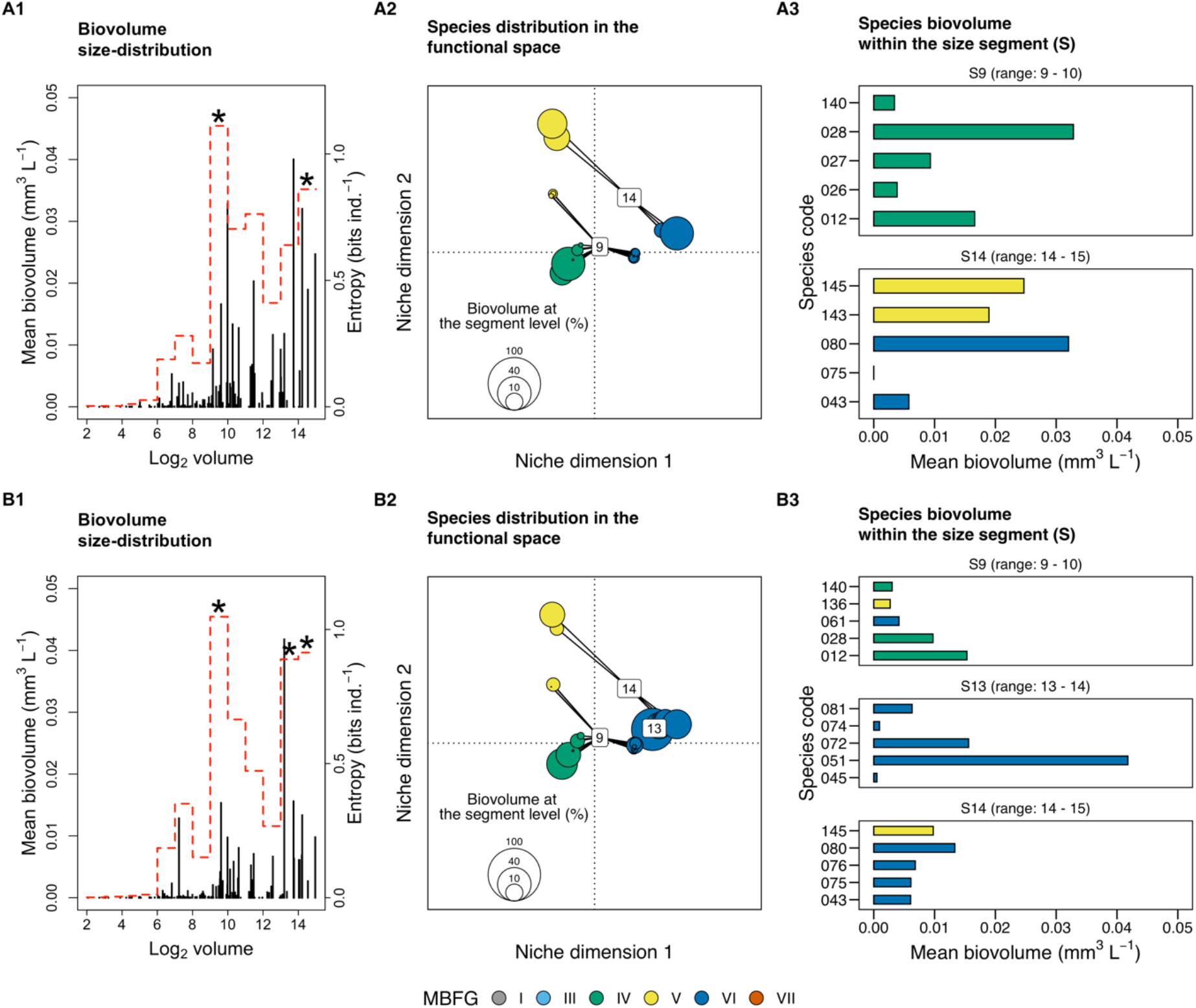
Seasonal distribution of phytoplankton biovolume along the body size axis, the ordination of species from the significant size segments (S) in the functional space, and the mean biovolume of the five most abundant species of each significant size segment during the (A) dry and (B) wet seasons of the Piabanha river, RJ. (1) Stem plots show size distribution in the sampling sites of the river Piabanha. Each stem represents a species with its body size (in log_2_) plotted on the abscissa and the mean biovolume plotted on the ordinate. The red dotted line indicates the entropy value of each size segment (i.e., unit of log_2_ volume) and the asterisk highlights the significant entropy values tested through 1000 randomizations. (2) The species of the corresponding significant size-segment are ordinated in the functional space. The size of the circles represents the species contribution to the total biovolume of the size segment, the black line connects species to the centroid (see equation 3), and the number in the centre of the clump indicates the size class it encompasses. (3) Bar plots show the biovolume of the five most abundant species from each significant size segment. Species are coloured according to their morphology-based functional groups (MBFG). The code for species can be found in the supplementary material, Table S1.

**Figure 5.**
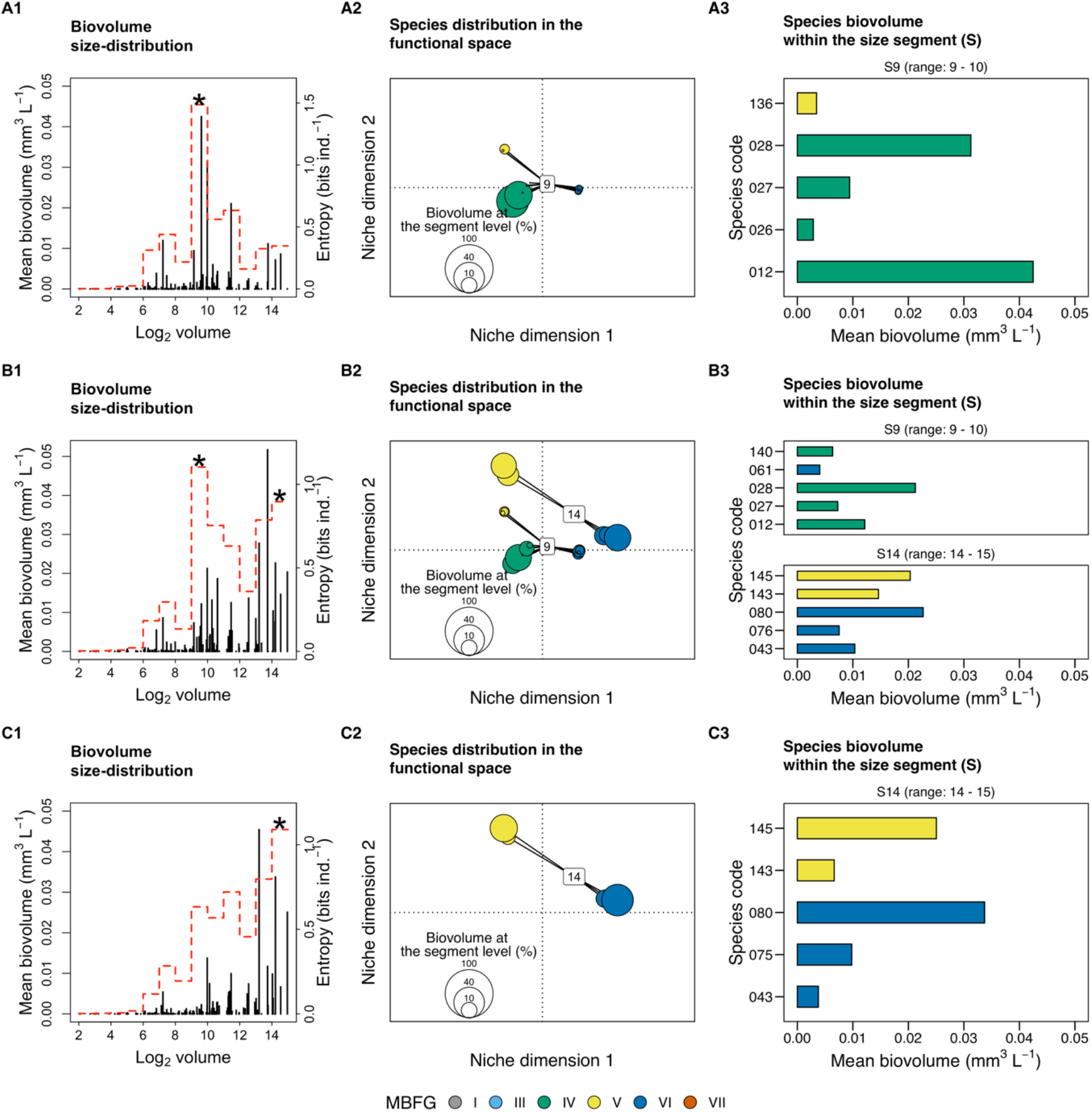
Spatial distribution of phytoplankton biovolume along the body size axis, the ordination of species from the significant size segments (S) in the functional space, and the mean biovolume of the five most abundant species of each significant size segment at the (A) upper, (B) medium, and (C) lower courses of the Piabanha river, RJ. (1) Stem plots show size distribution in the sampling sites of the river Piabanha. Each stem represents a species with its body size (in log_2_) plotted on the abscissa and the mean biovolume plotted on the ordinate. The red dotted line indicates the entropy value of each size segment (i.e., unit of log_2_ volume) and the asterisk highlights the significant entropy values tested through 1000 randomizations. (2) The species of the corresponding significant size-segment are ordinated in the functional space. The size of the circles represents the species contribution to the total biovolume of the size segment, the black line connects species to the centroid (see equation 3), and the number in the centre of the clump indicates the size class it encompasses. (3) Bar plots show the biovolume of the five most abundant species from each significant size segment. Species are coloured according to their morphology-based functional groups (MBFG). The code for species can be found in the supplementary material, Table S1.

Species from the same MBFG tended to cluster in the functional space even if they belonged to different clumps (Figure 4 – 5; Column 2). Yet, species within the same MBFG did not attain the highest abundance in more than one clump (Figure 4 – 5; Column 3). The mean biovolume of species within clumps differed between seasons, but the identity of the most abundant species did not vary (Figure 4 – Column 3). *Pseudanabaena* sp. (spp. 028) and *P. catenata* (spp. 012) had the highest biovolumes of clump I at both dry (Figure 4 – A2) and wet (Figure 4 – B3) seasons. Within clump II, *Synedra* sp. (spp. 080) attained the highest biovolume during the dry season (Figure 4 – A3) while *Cymbella* sp. (spp. 051) had the highest biovolume in the wet season (Figure 4 – B3).

Regarding the river stretches, only the clump I had significant entropy values for species from the S9 (Figure 5 – A1), with *Pseudanabaena* sp. (spp. 028) and *P. catenata* (spp. 012) contributing most of the biovolume (Figure 5 – A3). At the medium course, both clumps I and II had significant entropy values with *Pseudanabaena* sp. (spp. 028) attaining the highest biovolume within clump I, and *Synedra* sp. (spp. 080) attaining the highest biovolume within clump II (Figure 5 – C3). At the lower course, only clump II had significant entropy values at the S14 with *Synedra* sp. (spp. 080) as the most representative species (Figure 5 – C3).

Mantel tests showed that species’ pairwise differences in biovolume correlated with functional dissimilarity irrespectively of the season or river stretch when the whole community was analyzed (Table 3). The correlation was highest during the wet season (Mantel *r* = 0.23, p < 0.01) and at the lower course (Mantel *r* = 0.26, p < 0.01). For the clump-level pairwise differences, we only found a significant correlation at the upper course (Mantel *r* = 0.23, p < 0.02; Table 3). In contrast, functional distinctiveness at clump level presented a significant positive relationship for both the dry season (⍰ = 16.44, R^2^ = 0.40, p < 0.01) and the wet season (⍰ =18.32, R^2^ = 0.40, p < 0.01), and also for the upper (⍰ = 5.68, R^2^ = 0.40, p < 0.01) and medium (⍰ = 14.46, R^2^ = 0.34, p = 0.02) courses (Table 4), indicating that species with the most distinct trait combinations within the clumps also attain the highest biovolume. Essentially, such pattern was observed only for the species within clump I, except during the wet season where species from clump II also showed a significant positive relationship (⍰ = - 17.83, p = 0.02; Table 4)

**Table 3.** Mantel correlation results. Mantel correlation between the differences in species biovolume and functional dissimilarity for the whole community, and separately for the species within significant clumps (Figure 3 – 4) along the seasons (dry and wet) and river stretches (upper, medium, and lower courses). Species number of each stratum (whole community or clumps) are given. The relationships were tested for significance using 999 permutations, whenever possible.

| *Seasons and river stretches* | *Stratum* | *Number of species* | *Mantel r* | *p-value (permutations)* |
| --- | --- | --- | --- | --- |
| *Dry season* |  |  |  |  |
|  | Whole community | 135 | **0.25** | **< 0.01 (999)** |
|  | Clump I | 22 | 0.13 | 0.09 (999) |
|  | Clump II | 4 | -0.14 | 0.71 (23) |
| *Wet Season* |  |  |  |  |
|  | Whole community | 123 | **0.29** | **< 0.01 (999)** |
|  | Clump I | 20 | 0.06 | 0.24 (999) |
|  | Clump II | 11 | -0.16 | 0.69 (999) |
| *Upper course* |  |  |  |  |
|  | Whole community | 100 | **0.17** | **< 0.01 (999)** |
|  | Clump I | 17 | **0.23** | **0.02**<br>(999) |
| *Medium course* |  |  |  |  |
|  | Whole community | 135 | **0.21** | **< 0.01**<br>(999) |
|  | Clump I | 23 | 0.06 | 0.22<br>(999) |
|  | Clump II | 5 | -0.16 | 0.65<br>(119) |
| *Lower course* |  |  |  |  |
|  | Whole community | 122 | **0.33** | **<0.01**<br>(999) |
|  | Clump II | 5 | -0.23 | 0.75<br>(119) |
| *Note:* | | Bold values indicate significant correlations ($p < 0.05$). | | |

**Table 4.** Linear model results. Regression parameters of the relationship between species biovolume and functional distinctiveness at the clump level. The coefficients are shown along with the p-values of each independent variable.

| Dependent variable: $\log_{10}$ Biovolume | | | | | |
| --- | --- | --- | --- | --- | --- |
| Independent variables | Dry season | Wet season | Upper course | Medium course | Lower course |
| $F_{\text{Dist}}$ | 16.44<br>$p < 0.01$ | 18.33<br>$p < 0.01$ | 5.68<br>$p > 0.01$ | 14.46<br>$p = 0.02$ | 9.65<br>$p = 0.06$ |
| Clump II | 12.29<br>$p = 0.30$ | 17.73<br>$p = 0.01$ | | 11.61<br>$p = 0.29$ | 5.77<br>$p = 0.48$ |
| $F_{\text{Dist}} \times \text{Clump II}$ | -11.90<br>$p = 0.34$ | -17.83<br>$p = 0.02$ | | -11.12<br>$p = 0.34$ | -4.57<br>$p = 0.59$ |
| Intercept | -18.39<br>p < 0.01 | -20.420<br>p < 0.01 | -7.26<br>p < 0.01 | -16.64<br>p < 0.01 | -12.52<br>p = 0.01 |
| Observations | 26 | 31 | 17 | 28 | 26 |
| Adjusted R<sup>2</sup> | 0.41 | 0.39 | 0.40 | 0.34 | 0.55 |
| F Statistic | 6.75<br>(df = 3; 22) | 7.58<br>(df = 3; 27) | 11.87<br>(df = 1; 15) | 5.70<br>(df = 3; 24) | 11.01<br>(df = 3; 22) |

## Discussion

Present results showed that (i) the clumps in body size are a conspicuous feature of phytoplankton community structure in riverine systems across seasons and river stretches; (ii) species within clumps showed a random distribution of biomass concerning their pairwise functional dissimilarity, but not at the whole-community level; and (iii) species biovolume generally increases for species far apart from the centroid of multivariate trait space (i.e., functional distinctiveness) within clumps. Altogether these results support the Emergent neutrality hypothesis and show studying species beyond pairwise interactions help to explain the biomass distribution of functionally similar species, paving the way to analyze intra-clumps trait distributions.

Multimodal aggregation of species biovolume along body size axis only points to the integration of niche-based processes and neutrality driving community assembly (Vergnon et al. 2009), supporting H_1_. Alternative hypothesis such as pure neutrality (Hubbell 2001) or high dimensional hypothesis (HDH, Clark et al. 2007) are not supported by present results because pure neutrality predicts a uniform distribution of species biovolume and traits along the niche axis (Hubbell 2001), and the HDH does not predict any particular trait distribution (Vergnon et al. 2009, Ingram et al. 2018).

One alternative theory that is likely to explain clumpy aggregations is Holling’s textural hypothesis (Holling 1992), which suggests that multimodal species size distribution is the result of environmental constraints. Our results do not support textural hypothesis, as river stretches and seasons were markedly different in hydrology, nutrient concentrations, and other relevant descriptors of riverine landscapes fluxes, but that was not reflected in the stable clumpy size structure of the phytoplankton registered in the present study (Figure 4). The stability found in the clumps agree with empirical results registered in Segura et al. (2011, 2013b) and theoretical findings on the location of clumps (Fort et al. 2009). However, morphological trait composition of species in the different clumps reflected different environmental templates. The dominant species from the clump I belonged to MBFG IV (Figure 4–5) and presented highest biovolume under low-flow and high nutrient conditions (Figure 3), which is in line with previous findings for this MBFG (Chen et al. 2015). Within the second clump (II), the most abundant species belonged to MBFG VI and had their highest biovolume under high-flow and turbid conditions, in line with the ecology of silicious organisms (diatoms) able to cope with turbulent environments (Bortolini and Bueno 2017). The empirical evidence is consistent with studies from coastal and estuarine environments (Segura et al. 2011, 2013b) and are in line with recent modelling results suggesting that clumpy patterns arise in environments subjected to resources fluctuation (Sakavara et al. 2018), such as rivers. The trade-off between resources among competing species (Tilman 1982), which is a required ingredient for the emergence of clumps should be further explored.

The analysis of multiple trait dimensions, combining morphology based functional groups (MBFG) and quantitative distance metrics helped to describe the changes in species traits within clumps. We found that the species from the same clump are distributed across multiple MBFGs but only species from the same MBFG attain the highest biovolume within a clump, reinforcing that body size is a good proxy for niche differences of the species (Blanckenhorn 2000, Gallego et al. 2019). MBFGs helped to detect differences at a finer degree because they synthesize multiple trait dimensions as suggested previously to understand community organization (D’Andrea et al. 2018). Given that species within the same MBFG share similar ecological strategies (Kruk et al. 2010, Kruk and Segura 2012) under EN premises they should also perform similarly (Scheffer et al. 2018). This is in line with the significant functional dissimilarity observed at the whole community level but not for each clump separately, agreeing with H_2_. The effects of traits in species’ fitness are context-dependent, however, whereas the use of traits for assigning MBFGs is static. Testing the significance of traits given the observed environmental conditions might help to unveil community assembling processes at an even finer degree (Kremer et al. 2017).

We also outlined the role of functional similarity in community assembly by studying the effects of functional distinctiveness on species biovolume at clump level. Within the clump I, species biovolume increased with functional distinctiveness, but this pattern was weaker within clump II (Table 4). It may be that such patterns stem from the fact that phytoplankton growth-rate decreases with body size while increases with surface volume ratio, providing that populations of smaller species (such as in clump I) are less sensitive to losses by flushing rates (Kruk et al. 2010). On the other hand, large-sized species have often an elongated shape that provides advantage under turbulent conditions with low light availability (Reynolds et al. 1994). Differently, aerotopes and mucilage are useful to reach the surface in deep stratified lakes but are not key in small rivers or streams where turbulent fluxes dominates and these traits are not useful to recuperate the position in the water column. Furthermore, in rivers, large-bodied phytoplanktonic species are often randomly introduced from different habitats (e.g. periphyton or epiphyton) (Wang et al. 2014, Descy et al. 2017), which has also been found true for the Piabanha river especially under high flow conditions (Graco-Roza et al. 2020). This mechanism can help to explain the weak relationship between functional distinctiveness and biovolume within clump II, which explains the niche overlap found in large-sized species as the result of immigration and emigration out of the pelagic zone.

Emergent neutrality results from eco-evolutionary processes that lead species selection towards a limited number of functional groups (Scheffer and van Nes 2006). This implies that the clumps observed here are not likely a result of competitive exclusion at the Piabanha river, but a convergent evolution of competing species over time (MacArthur and Levins 1967). Therefore, even when the competition rates are relaxed due to sufficient nutrient supply, some other limiting factors that are not consumed by biotic organisms such as heat energy or turbulence determine species biovolume. Looking at species differences at a high-order level (instead of pairwise differences) helped to detect the effects of trait composition on the biovolume distribution in quasi-neutral clumps. In fact, our results showed that it is possible to predict the biovolume of species within clumps, but only when immigration from different habitats are relaxed and biotic interactions are more likely to occur. Therefore, our findings partially agree with H_3_-there is a positive relationship between species abundance and species functional distinctiveness within clumps, but the environmental conditions seem to play a key role in the outcome.

There are some possible influential aspects in our study design that should be discussed. First, given that phytoplankton communities have short generation time (Reynolds 2006), the monthly resolution makes difficult to capture the turnover in species abundance rank at the finest possible scale. Studying the daily or even weekly variation in abundance rank would help us to disentangle the so-called stochastic abundance fluctuations (Caracciolo et al. 2021). However, it has been shown that the study of phytoplankton community processes on monthly to yearly time scales helps to understand the long-term ecological and evolutive dynamics of communities (Segura et al. 2011). Secondly, considering intraspecific trait variability allows one to disentangle species responses from environmental variations (Wong and Carmona 2021). Here, we assessed traits at species level and comparing the overlap in trait values between species might unveil the quasi-neutral relationship between pairs or clumps of species.

In summary, we provided evidence of both neutral and niche mechanisms driving planktonic community assembly and support the view that Emergent neutrality is a likely mechanism to explain species coexistence in an open and environmentally heterogeneous ecosystem. The use of MBFG classification and functional space to describe species within clumps revealed that under the same size range, species with a greater degree of functional similarity unpredictably alternate their dominance. The position and dominance of the clumps were related to the environmental conditions, but the biovolume of species within the clumps was better predicted by functional distinctiveness than by pairwise functional similarity. This addresses the difficulty to avoid the ghost of hidden niches (Barabás et al. 2013) and also provides evidence from multiple angles that point to EN as a plausible mechanism in shaping species coexistence in riverine landscapes.

## Data accessibility

Data are available online: https://doi.org/10.5281/zenodo.4778444

## Supplementary material

R code used in the analysis: https://github.com/graco-roza/clumpy-coexistence-phytoplankton

## Acknowledgements

CGR PhD scholarship was funded by Fudação de Apoio a Pesquisa do Estado do Rio de Janeiro (FAPERJ), by Coordenação de Aperfeiçoamento de Pessoal de Nível Superior (CAPES), and by Ella and Georg Ehrnrooth foundation. MMM was partially supported by CNPq (303572/2017-5). Version 6 of this preprint has been peerreviewed and recommended by Peer Community In Ecology (https://doi.org/10.24072/pci.ecology.100083)

## Conflict of interest disclosure

The authors of this preprint declare that they have no financial conflict of interest with the content of this article.

## Appendix

Supplemental table 1: https://www.biorxiv.org/content/biorxiv/early/2021/05/05/869966/DC1/embed/media-1.pdf

## Notes

https://doi.org/10.5281/zenodo.4778444

### Competing Interest Statement

The authors have declared no competing interest.

### Summary of Updates

"Version 6 of this preprint has been peer-reviewed and recommended by Peer Community In Ecology (https://doi.org/10.24072/pci.ecology.100083)"

