## Supplemental Table 1 for "Clumpy coexistence in phytoplankton: The role of functional similarity in community assembly" (from another version of this preprint: version 5, 2021-05-05)

**Appendix 1** Clumpy coexistence in phytoplankton: The role of functional similarity in community assembly.

Table S1. Mophology based functional group (MBFG), individual volume (V), surface volume ratio (SV), maximum linear dimension (MLD), presence of flagella (F), siliceous exoskeleton (Si), aerotopes (A), and the code used to describe each species observed during the study.

| Taxa | MBFG | V (µm³) | SV | MLD (µm) | F | Si | M | A | code |
|---|---|---|---|---|---|---|---|---|---|
| | | | | Cyanobacteria | | | | | |
| Chroococcales sp. | I | 4.2 | 3.0 | 2.0 | 0 | 0 | 0 | 0 | spp. 20 |
| Chroococcales sp. 2 | VII | 169.7 | 2.0 | 15.0 | 0 | 0 | 1 | 0 | spp. 21 |
| Chroococcales sp. 3 | VII | 171.6 | 1.9 | 13.0 | 0 | 0 | 1 | 0 | spp. 22 |
| Chroococcales sp. 4 | VII | 83.8 | 3.0 | 15.0 | 0 | 0 | 1 | 0 | spp. 23 |
| *Chroococcus* sp. | I | 22.5 | 1.7 | 3.5 | 0 | 0 | 0 | 0 | spp. 18 |
| *cf.Cyanobium* sp. | I | 8.4 | 2.6 | 4.0 | 0 | 0 | 0 | 0 | spp. 19 |
| *Geitlerinema* sp. | IV | 79.2 | 3.4 | 70.0 | 0 | 0 | 0 | 0 | spp. 1 |
| *Geitlerinema* sp. 2 | IV | 212.1 | 2.7 | 120.0 | 0 | 0 | 0 | 0 | spp. 2 |
| *Geitlerinema* sp. 3 | IV | 57.7 | 3.1 | 43.5 | 0 | 0 | 0 | 0 | spp. 3 |
| *Jaaginema* sp. | IV | 84.3 | 3.7 | 88.7 | 0 | 0 | 0 | 0 | spp. 5 |
| *Komvophoron* sp. | IV | 31.8 | 2.8 | 19.0 | 0 | 0 | 0 | 0 | spp. 4 |
| *Limnothrix* sp. | III | 1570.8 | 1.0 | 125.0 | 0 | 0 | 0 | 1 | spp. 6 |
| *Limnothrix* sp. 2 | III | 1256.6 | 1.0 | 100.0 | 0 | 0 | 0 | 1 | spp. 7 |
| *Limnothrix* sp. 3 | III | 2428.9 | 0.9 | 140.0 | 0 | 0 | 0 | 1 | spp. 8 |
| *Merismopedia* sp. | VII | 343.0 | 0.9 | 20.0 | 0 | 0 | 1 | 0 | spp. 24 |
| *Oscillatoria* sp. | IV | 1361.9 | 1.2 | 150.0 | 0 | 0 | 0 | 0 | spp. 9 |
| *Oscillatoria* sp. 2 | IV | 301.7 | 1.5 | 49.0 | 0 | 0 | 0 | 0 | spp. 10 |
| Oscillatoriales sp. | IV | 282.7 | 1.4 | 40.0 | 0 | 0 | 0 | 0 | spp. 33 |
| Oscillatoriales sp. 2 | IV | 141.4 | 1.4 | 20.0 | 0 | 0 | 0 | 0 | spp. 34 |
| Oscillatoriales sp. 4 | IV | 184.5 | 1.4 | 26.1 | 0 | 0 | 0 | 0 | spp. 35 |
| Oscillatoriales sp. 6 | IV | 85.0 | 1.7 | 16.0 | 0 | 0 | 0 | 0 | spp. 36 |
| *Pseudanabaena catenata* | IV | 785.4 | 2.0 | 250.0 | 0 | 0 | 0 | 0 | spp. 12 |
| *Pseudanabaena limnetica* | IV | 113.1 | 2.7 | 64.0 | 0 | 0 | 0 | 0 | spp. 11 |
| *Pseudanabaena* sp. | IV | 2828.6 | 1.2 | 294.0 | 0 | 0 | 0 | 0 | spp. 25 |

| | | | | | | | | | |
|---|---|---|---|---|---|---|---|---|---|
| *Pseudanabaena* sp. 2 | IV | 981.8 | 1.6 | 200.0 | 0 | 0 | 0 | 0 | spp. 26 |
| *Pseudanabaena* sp. 3 | IV | 568.6 | 2.0 | 181.0 | 0 | 0 | 0 | 0 | spp. 27 |
| *Pseudanabaena* sp. 4 | IV | 1011.2 | 1.6 | 206.0 | 0 | 0 | 0 | 0 | spp. 28 |
| *Pseudanabaena* sp. 5 | IV | 213.6 | 2.0 | 68.0 | 0 | 0 | 0 | 0 | spp. 29 |
| Pseudanabaenaceae sp. | IV | 176.7 | 2.7 | 100.0 | 0 | 0 | 0 | 0 | spp. 30 |
| Pseudanabaenaceae sp. 2 | IV | 106.2 | 3.1 | 80.0 | 0 | 0 | 0 | 0 | spp. 31 |
| *cf. Raphidiopsis* | IV | 254.5 | 2.2 | 100.0 | 0 | 0 | 0 | 0 | spp. 32 |
| Pseudanabaenaceae sp. 7 | IV | 21.2 | 4.1 | 27.0 | 0 | 0 | 0 | 0 | spp. 13 |
| *Romeria* sp. | IV | 125.7 | 2.1 | 40.0 | 0 | 0 | 0 | 0 | spp. 14 |
| *cf. Spirulina* | IV | 31.2 | 3.5 | 30.0 | 0 | 0 | 0 | 0 | spp. 15 |
| *Synechococcus* sp. | I | 4.2 | 3.0 | 4.0 | 0 | 0 | 0 | 0 | spp. 17 |
| *Synechocystis aquatilis* | IV | 32.3 | 1.5 | 4.0 | 0 | 0 | 0 | 0 | spp. 16 |
| Bacillariophyceae | | | | | | | | | |
| *Achnanthes cf. rupestoides* | VI | 569.4 | 0.9 | 29.0 | 0 | 1 | 0 | 0 | spp. 46 |
| *Achnanthes inflata* | VI | 8812.2 | 0.3 | 68.0 | 0 | 1 | 0 | 0 | spp. 45 |
| *Achnanthes* sp. | VI | 160.2 | 1.3 | 17.0 | 0 | 1 | 0 | 0 | spp. 47 |
| *Achnanthes* sp. 2 | VI | 99.0 | 1.8 | 21.0 | 0 | 1 | 0 | 0 | spp. 48 |
| *cf. Achnanthidium* sp. | VI | 48.7 | 3.1 | 31.0 | 0 | 1 | 0 | 0 | spp. 49 |
| *Amphora* sp. | VI | 340.9 | 1.1 | 30.0 | 0 | 1 | 0 | 0 | spp. 50 |
| *Aulacoseira granulata* | VI | 1263.6 | 0.7 | 49.0 | 0 | 1 | 0 | 0 | spp. 37 |
| *Aulacoseira* sp. | VI | 5657.2 | 0.6 | 147.0 | 0 | 1 | 0 | 0 | spp. 38 |
| *Aulacoseira* sp.2 | VI | 4834.9 | 0.5 | 76.0 | 0 | 1 | 0 | 0 | spp. 39 |
| Centrales sp. | VI | 5195.4 | 0.6 | 135.0 | 0 | 1 | 0 | 0 | spp. 44 |
| *Cyclotella cf. meneghiniana* | VI | 716.8 | 0.6 | 15.0 | 0 | 1 | 0 | 0 | spp. 40 |
| *Cyclotella* sp. | VI | 410.5 | 0.8 | 12.0 | 0 | 1 | 0 | 0 | spp. 41 |
| *Cyclotella* sp. 2 | VI | 190.1 | 1.0 | 8.0 | 0 | 1 | 0 | 0 | spp. 42 |
| *Cymbela* sp. | VI | 9433.0 | 0.3 | 64.0 | 0 | 1 | 0 | 0 | spp. 51 |
| *cf. Encyonema* sp. | VI | 226.2 | 1.2 | 18.0 | 0 | 1 | 0 | 0 | spp. 52 |
| *Fragillaria capucina* | VI | 648.0 | 0.9 | 33.0 | 0 | 1 | 0 | 0 | spp. 53 |
| *Fragillaria capucina var. fragilarioides* | VI | 989.6 | 0.7 | 36.0 | 0 | 1 | 0 | 0 | spp. 55 |
| *Fragillaria capucina var. gracilis* | VI | 730.4 | 0.8 | 31.0 | 0 | 1 | 0 | 0 | spp. 54 |
| *Fragillaria* sp. | VI | 487.0 | 1.0 | 31.0 | 0 | 1 | 0 | 0 | spp. 56 |
| *cf. Frustulia* sp. | VI | 1526.8 | 0.5 | 27.0 | 0 | 1 | 0 | 0 | spp. 57 |
| *Gomphonema augur* | VI | 78.6 | 1.6 | 27.0 | 0 | 1 | 0 | 0 | spp. 58 |
| *Gomphonema cf. olivaceum* | VI | 150.1 | 1.2 | 31.0 | 0 | 1 | 0 | 0 | spp. 59 |

| | | | | | | | | | |
|---|---|---|---|---|---|---|---|---|---|
| *Gomphonema cf. truncatum* | VI | 139.9 | 1.3 | 33.0 | 0 | 1 | 0 | 0 | spp. 60 |
| *Gomphonema* sp. | VI | 758.7 | 0.7 | 23.0 | 0 | 1 | 0 | 0 | spp. 61 |
| *Gomphonema* sp. 2 | VI | 212.1 | 1.2 | 18.0 | 0 | 1 | 0 | 0 | spp. 62 |
| *Gyrosigma/Pleurosigma* | VI | 8278.1 | 0.3 | 61.9 | 0 | 1 | 0 | 0 | spp. 63 |
| *Hantszchia* sp. | VI | 966.0 | 1.0 | 46.0 | 0 | 1 | 0 | 0 | spp. 64 |
| *Melosira* sp. | VI | 17241.1 | 0.2 | 28.0 | 0 | 1 | 0 | 0 | spp. 43 |
| Navicula sp. | VI | 2660.9 | 0.5 | 44.0 | 0 | 1 | 0 | 0 | spp. 65 |
| *Navicula* sp. 2 | VI | 546.6 | 0.9 | 28.6 | 0 | 1 | 0 | 0 | spp. 66 |
| *Navicula* sp. 3 | VI | 1286.5 | 0.6 | 25.9 | 0 | 1 | 0 | 0 | spp. 67 |
| *Navicula* sp. 4 | VI | 1060.3 | 0.6 | 25.0 | 0 | 1 | 0 | 0 | spp. 68 |
| *Naviculales* | VI | 2937.4 | 0.4 | 34.3 | 0 | 1 | 0 | 0 | spp. 69 |
| Nitzschia sp. | VI | 395.8 | 1.2 | 41.7 | 0 | 1 | 0 | 0 | spp. 70 |
| *Nitzschia* sp. 2 | VI | 99.0 | 1.8 | 21.0 | 0 | 1 | 0 | 0 | spp. 71 |
| Pennales sp. | VI | 2660.9 | 0.5 | 43.8 | 0 | 1 | 0 | 0 | spp. 82 |
| Pennales sp. 10 | VI | 754.0 | 0.7 | 19.6 | 0 | 1 | 0 | 0 | spp. 90 |
| Pennales sp. 11 | VI | 94.3 | 1.3 | 8.0 | 0 | 1 | 0 | 0 | spp. 91 |
| Pennales sp. 12 | VI | 1005.3 | 0.6 | 20.0 | 0 | 1 | 0 | 0 | spp. 92 |
| Pennales sp. 13 | VI | 824.7 | 0.8 | 29.7 | 0 | 1 | 0 | 0 | spp. 93 |
| Pennales sp. 2 | VI | 2544.7 | 0.5 | 40.0 | 0 | 1 | 0 | 0 | spp. 83 |
| Pennales sp. 3 | VI | 1237.0 | 0.6 | 24.7 | 0 | 1 | 0 | 0 | spp. 84 |
| Pennales sp. 4 | VI | 2353.8 | 0.5 | 37.0 | 0 | 1 | 0 | 0 | spp. 85 |
| Pennales sp. 5 | VI | 1357.2 | 0.6 | 27.0 | 0 | 1 | 0 | 0 | spp. 86 |
| Pennales sp. 6 | VI | 7861.8 | 0.3 | 55.0 | 0 | 1 | 0 | 0 | spp. 87 |
| Pennales sp. 7 | VI | 1696.5 | 0.5 | 30.0 | 0 | 1 | 0 | 0 | spp. 88 |
| Pennales sp. 8 | VI | 348.7 | 1.2 | 37.0 | 0 | 1 | 0 | 0 | spp. 89 |
| *Pinnularia* sp. | VI | 13589.6 | 0.3 | 71.0 | 0 | 1 | 0 | 0 | spp. 72 |
| *Pinnularia* sp. 2 | VI | 6013.0 | 0.4 | 49.3 | 0 | 1 | 0 | 0 | spp. 73 |
| *Pinnularia* sp. 7 | VI | 18746.1 | 0.3 | 62.3 | 0 | 1 | 0 | 0 | spp. 76 |
| *Pinullaria* sp. 4 | VI | 10449.1 | 0.3 | 48.7 | 0 | 1 | 0 | 0 | spp. 74 |
| *Pinullaria* sp. 6 | VI | 16760.3 | 0.3 | 64.4 | 0 | 1 | 0 | 0 | spp. 75 |
| *Stauroneis borrichii* | VI | 659.7 | 0.8 | 21.0 | 0 | 1 | 0 | 0 | spp. 77 |
| *Surirella cf. minuta* | VI | 1520.5 | 0.5 | 16.0 | 0 | 1 | 0 | 0 | spp. 78 |
| *Surirella* sp. | VI | 4002.4 | 0.4 | 26.0 | 0 | 1 | 0 | 0 | spp. 79 |
| *Synedra* sp. | VI | 19045.7 | 0.4 | 160.9 | 0 | 1 | 0 | 0 | spp. 80 |
| *cf. Urosolenia* sp. | VI | 13845.8 | 0.3 | 61.0 | 0 | 1 | 0 | 0 | spp. 81 |
| Chlorophyceae | | | | | | | | | |

| | | | | | | | | | |
|---|---|---|---|---|---|---|---|---|---|
| *Actinastrum cf. hantzchii* | IV | 388.8 | 1.2 | 25.0 | 0 | 0 | 0 | 0 | spp. 95 |
| *Actinastrum* sp. | IV | 115.2 | 1.9 | 16.0 | 0 | 0 | 0 | 0 | spp. 94 |
| *Ankistrodesmus* sp. | IV | 578.6 | 2.0 | 37.5 | 0 | 0 | 0 | 0 | spp. 96 |
| *Chlorella* sp. 1 | IV | 65.5 | 1.2 | 5.0 | 0 | 0 | 0 | 0 | spp. 97 |
| *Chlorella* sp. 2 | IV | 46.8 | 1.3 | 4.5 | 0 | 0 | 0 | 0 | spp. 98 |
| *Chlorella* sp. 3 | IV | 87.1 | 1.1 | 5.5 | 0 | 0 | 0 | 0 | spp. 99 |
| Chlorococcales sp. | IV | 69.5 | 1.2 | 5.1 | 0 | 0 | 0 | 0 | spp. 127 |
| Chlorococcales sp. 2 | IV | 212.2 | 0.8 | 7.4 | 0 | 0 | 0 | 0 | spp. 128 |
| Chlorococcales sp. 3 | IV | 33.5 | 1.5 | 4.0 | 0 | 0 | 0 | 0 | spp. 129 |
| Chlorococcales sp. 4 | IV | 47.4 | 1.3 | 4.5 | 0 | 0 | 0 | 0 | spp. 130 |
| Chlorococcales sp. 5 | VII | 235.6 | 1.1 | 51.2 | 0 | 0 | 1 | 0 | spp. 131 |
| Chlorococcales sp. 6 | VII | 2822.2 | 0.8 | 15.0 | 0 | 0 | 1 | 0 | spp. 132 |
| Chlorococcales sp. 7 | VII | 293.2 | 1.3 | 17.0 | 0 | 0 | 1 | 0 | spp. 133 |
| *Coelastrum* sp. | IV | 8181.2 | 0.2 | 25.0 | 0 | 0 | 0 | 0 | spp. 100 |
| *Crucigenia fenestrata* | IV | 217.8 | 1.0 | 6.1 | 0 | 0 | 0 | 0 | spp. 101 |
| *Crucigenia* sp. 2 | IV | 198.0 | 1.0 | 5.5 | 0 | 0 | 0 | 0 | spp. 102 |
| *Crucigeniella* sp. | IV | 1139.0 | 0.6 | 13.6 | 0 | 0 | 0 | 0 | spp. 103 |
| *Desmodesmus quadricauda* | IV | 188.5 | 1.6 | 9.6 | 0 | 0 | 0 | 0 | spp. 104 |
| *Desmodesmus* sp. 1 | IV | 301.6 | 1.3 | 8.6 | 0 | 0 | 0 | 0 | spp. 105 |
| *Desmodesmus* sp. 2 | IV | 469.1 | 1.2 | 14.4 | 0 | 0 | 0 | 0 | spp. 106 |
| *Desmodesmus* sp. 3 | IV | 113.1 | 1.7 | 5.9 | 0 | 0 | 0 | 0 | spp. 107 |
| *Desmodesmus* sp. 4 | IV | 94.3 | 1.8 | 5.0 | 0 | 0 | 0 | 0 | spp. 108 |
| *Desmodesmus* sp. 5 | IV | 418.9 | 1.1 | 8.0 | 0 | 0 | 0 | 0 | spp. 109 |
| *Dictyosphaerium* sp. | VII | 112.3 | 1.7 | 3.5 | 0 | 0 | 1 | 0 | spp. 110 |
| *cf. Koliella* sp. | I | 6.5 | 2.8 | 4.0 | 0 | 0 | 0 | 0 | spp. 111 |
| *Monoraphidium arcuatum* | I | 14.8 | 3.4 | 17.5 | 0 | 0 | 0 | 0 | spp. 112 |
| *Monoraphidium contortum* | I | 7.7 | 4.6 | 17.5 | 0 | 0 | 0 | 0 | spp. 113 |
| *Monoraphidium convolutum* | I | 9.3 | 3.4 | 11.0 | 0 | 0 | 0 | 0 | spp. 114 |
| *Monoraphidium komarkovae* | IV | 23.6 | 4.0 | 40.0 | 0 | 0 | 0 | 0 | spp. 115 |
| *Monoraphidium minutum* | I | 18.9 | 2.1 | 8.0 | 0 | 0 | 0 | 0 | spp. 116 |
| *Oocystis* sp. | IV | 384.9 | 0.7 | 15.0 | 0 | 0 | 0 | 0 | spp. 117 |
| *Oocystis* sp. 2 | IV | 150.8 | 0.9 | 8.2 | 0 | 0 | 0 | 0 | spp. 118 |
| *Oocystis* sp. 3 | IV | 1140.4 | 0.5 | 18.3 | 0 | 0 | 0 | 0 | spp. 119 |
| *Pandorina* sp. | IV | 1415.8 | 0.4 | 15.7 | 0 | 0 | 0 | 0 | spp. 120 |
| *Pediastrum* sp. | IV | 5717.7 | 0.9 | 56.0 | 0 | 0 | 0 | 0 | spp. 121 |

| | | | | | | | | | |
|---|---|---|---|---|---|---|---|---|---|
| *Pediastrum* sp. 2 | IV | 3518.6 | 1.3 | 50.0 | 0 | 0 | 0 | 0 | spp. 122 |
| *Scenedesmus* sp. | IV | 301.6 | 1.3 | 8.9 | 0 | 0 | 0 | 0 | spp. 123 |
| *Scenedesmus* sp. 2 | IV | 234.6 | 1.2 | 13.5 | 0 | 0 | 0 | 0 | spp. 124 |
| *Scenedesmus* sp. 3 | IV | 829.4 | 0.9 | 10.6 | 0 | 0 | 0 | 0 | spp. 125 |
| Scenedesmus sp. 4 | IV | 50.3 | 2.5 | 6.0 | 0 | 0 | 0 | 0 | spp. 126 |
| Cryptophyceae | | | | | | | | | |
| *cf. Cryptomonas* sp. | V | 785.4 | 0.5 | 14.7 | 1 | 0 | 0 | 0 | spp. 134 |
| *cf. Cryptomonas* sp. 2 | V | 551.4 | 0.6 | 12.8 | 1 | 0 | 0 | 0 | spp. 135 |
| Cryptophyceae sp. | V | 837.8 | 0.5 | 16.0 | 1 | 0 | 0 | 0 | spp. 136 |
| Cryptophyceae sp. 2 | V | 368.6 | 0.7 | 11.0 | 1 | 0 | 0 | 0 | spp. 137 |
| Cryptophyceae sp. 3 | V | 188.5 | 0.9 | 10.0 | 1 | 0 | 0 | 0 | spp. 138 |
| Cryptophyceae sp. 4 | V | 31.56 | 1.54 | 3.9 | 1 | 0 | 0 | 0 | spp. 149 |
| Cryptophyceae sp. 5 | V | 77.95 | 1.13 | 5.3 | 1 | 0 | 0 | 0 | spp. 150 |
| Zygnematophyceae | | | | | | | | | |
| *Cosmarium* sp. | IV | 1256.6 | 0.6 | 12.0 | 0 | 0 | 0 | 0 | spp. 139 |
| *Staurastrum* sp. | IV | 751.4 | 1.0 | 14.2 | 0 | 0 | 0 | 0 | spp. 140 |
| *Staurodesmus* sp. | IV | 1118.4 | 0.8 | 35.0 | 0 | 0 | 0 | 0 | spp. 141 |
| Chrysophyceae | | | | | | | | | |
| *Dinobryon* sp. | IV | 815.0 | 0.7 | 38.0 | 0 | 0 | 0 | 0 | spp. 142 |
| Euglenophyceae | | | | | | | | | |
| *Euglena* sp. 1 | V | 2508.1 | 0.3 | 30.5 | 1 | 0 | 0 | 0 | spp. 143 |
| *Euglena* sp. 2 | V | 3951.3 | 0.8 | 42.6 | 1 | 0 | 0 | 0 | spp. 144 |
| *Euglena* sp. 3 | V | 523.6 | 0.6 | 10.0 | 1 | 0 | 0 | 0 | spp. 145 |
| *Phacus* sp. | V | 1436.8 | 0.4 | 14.0 | 1 | 0 | 0 | 0 | spp. 146 |
| *Strombomonas* sp. | V | 31.1 | 1.5 | 3.9 | 1 | 0 | 0 | 0 | spp. 147 |
| *Trachelomonas* sp. | V | 78.0 | 1.1 | 5.3 | 1 | 0 | 0 | 0 | spp. 148 |
